# A piRNA modulates the levels of 20-hydroxyecdysone in the ovary of the German cockroach

**DOI:** 10.1101/2024.10.10.617606

**Authors:** Judit Gonzalvo, Nuria Farrus, Jorge Escudero, Petra Berková, Martin Moos, Marcela Nouzova, Fernando G. Noriega, David Pujal, Josep Bau, Maria-Dolors Piulachs

## Abstract

PIWI-interacting RNAs (piRNAs) are small non-coding RNAs, typically 26 to 31 nucleotides long, originally known for silencing transposable elements (TEs), thus maintaining genomic stability. However, recent research has revealed additional regulatory roles. In this study, we investigate piRNA-305221, which is highly expressed in the ovaries of the German cockroach, *Blattella germanica*, to understand its involvement in oogenesis and reproduction. piRNA-305221 is found in germinal and somatic cells during the gonadotropic cycle, and is maternally provided to the egg. Its expression correlates with critical ovarian events, such as endoreplication and follicular cell differentiation, suggesting regulatory functions beyond TE silencing. Functional knockdown using antisense oligonucleotides (ASOs) resulted in delayed oviposition, malformed oothecae, and reduced offspring viability. Gene expression analysis revealed that the reduction of piRNA-305221 decreased *shade* mRNA levels, impairing the conversion of ecdysone to its active form, 20-hydroxyecdysone, and a concomitant increase in expression of upstream steroidogenic genes (*spook, phantom, disembodied*). These results indicate that piRNA-305221 may regulate steroidogenesis through direct or indirect control of mRNA targets. This study highlights the broader regulatory functions of piRNAs and demonstrates the utility of ASO-mediated knockdown in functional studies of non-coding RNAs.

## Introduction

The piwi-interacting RNAs (piRNAs) are small non-coding RNAs (sncRNAs) ranging from 26 to 31 nt in length, whose biogenesis is associated with PIWI proteins subfamily (Hirakata and Siomi, 2016; Le Thomas et al., 2014). In terms of sequence, piRNAs are poorly conserved even among closely related species, suggesting their generation and function is highly species-specific (Aravin et al., 2007; O’Donnell and Boeke, 2007; Wang et al., 2019). In the fly *Drosophila melanogaster*, two pathways of piRNA biogenesis have been described: the primary pathway active in somatic cells and the secondary, or “ping-pong” pathway, which operates in germ cells. These pathways silence transposable elements (TEs) through transcriptional and post-transcriptional mechanisms (Gleason et al., 2018; Hirakata and Siomi, 2016). However, evidence from other arthropods indicates that both piRNA pathways can function in somatic as well as germinal tissues (Cerqueira de Araujo et al., 2022; Yamashita et al., 2024).

The first function associated with piRNAs was the repression of TEs, as part of an evolutionarily conserved mechanism that ensures genomic stability in germ cells (Aravin et al., 2007; Senti et al., 2015). However, recent studies have shown that piRNAs can also originate from coding sequences, intergenic regions unrelated to TEs, and 5’ and 3’ untranslated regions (UTRs) of mRNAs, which suggests that piRNAs regulate gene expression (Iki et al., 2023; Jensen et al., 2020; Lambert et al., 2019; Rojas-Ríos et al., 2018). In this context, piRNAs have been implicated in diverse functions, including embryogenesis, germ cell specification, primary sex determination, and the establishment of epigenetic states (Gleason et al., 2018; Gou et al., 2014; Iki et al., 2023; Kiuchi et al., 2014).

In a previous study on the German cockroach, *Blattella germanica*, we identified a significant number of piRNAs maternally provided to the zygote, which are abundant in early embryo stages (Belles et al., 2024; Llonga et al., 2018). In the present work, we approached the functional study of one of these piRNAs by reducing its expression using an antisense oligonucleotide (ASO). ASOs are short (18-30 nt) single-stranded DNA molecules that bind specifically to RNA to modulate gene expression (Roberts et al., 2020). Since the late 1970s, ASOs have been used as therapeutics against different diseases, as they can be designed with high specificity against their mRNA targets (Crooke et al., 2021). ASOs bind to the corresponding mRNA target by Watson–Crick base pairing, forming RNA-DNA heteroduplexes that recruit RNAse-H endonuclease, leading to cleavage and degradation of the target RNA. Despite their proven potential, ASOs have been rarely used to regulate insect gene expression (Oberemok et al., 2018).

*B. germanica* has a panoistic ovary type, in which all germline stem cells are converted into functional oocytes, with transcriptionally active nuclei (Bu□ning, 1994). During oogenesis, only the basal oocyte of each ovariole matures in every gonadotropic cycle, while the rest of the oocytes present in the vitellarium remain arrested until the next cycle begins (Irles and Piulachs, 2014; Rumbo et al., 2023; Belles et al., 2024). Once released from the germarium, the oocyte is surrounded by a monolayer of follicular cells (FCs), thus establishing an ovarian follicle (Rumbo et al., 2023). The FCs mature and change their characteristics in parallel to oocyte growth. In the sixth (last) nymphal instar, when the basal ovarian follicle (BOF) begins to mature, the FCs proliferate, increasing their number until they completely cover the oocyte (Irles and Piulachs, 2014; Belles et al., 2024). Later, in 3-day-old adults, cytokinesis in FCs is arrested, and cells become binucleated. Under the control of the juvenile hormone circulating in the hemolymph, the FCs contract their cytoplasm, thus leaving large intercellular spaces (Davey, 1981; Davey and Huebner, 1974), which facilitate the access of vitellogenin to the oocyte membrane. Henceforth, the oocytes in the BOFs grow exponentially. At the end of the gonadotropic cycle, the FCs close the intercellular spaces definitively (Belles et al., 2024) and chorion synthesis begins, triggered by the ecdysone production in the ovary (Belles et al., 1993; Pascual et al., 1992; Belles et al., 2024), a process that takes just a few hours (Irles et al., 2009b).

Several genes involved in ecdysone biosynthesis have been characterized in the ovaries of *B. germanica*, namely *neverland* (*nvd*), *spook* (spo), *spookiest* (*spot*), *phantom* (*phm*), *disembodied* (*dib*), and *shadow* (*sad*) (Belles et al., 2024). s*hade* (*shd*) catalyzes the conversion of ecdysone into its biologically active form, 20-hydroxyecdysone (20E) (Petryk et al., 2003). Among other transducers, the action of ecdysone is mediated by *E75*, an early gene in the ecdysone signaling cascade, and *fushi tarazu-f1* (*ftz-f1*), a late gene. *ftz-f1* also regulates the expression of steroidogenic genes and helps to maintain the correct cytoskeleton organization in the BOF at the end of the gonadotropic cycle (Alborzi and Piulachs, 2023; Farrus et al., 2024). When choriogenesis is complete, the cytoskeleton of the follicular epithelium in the BOFs rearranges, facilitating the release of the egg into the oviduct. (Alborzi and Piulachs, 2023; Belles et al., 2024). The eggs are then oviposited into an egg case or ootheca. In *B. germanica*, the female carries the ootheca attached to the genital atrium throughout embryogenesis, which lasts 18 days under our laboratory conditions (Piulachs et al., 2010; Belles et al., 2024).

## 2. Materials and methods

### 2.1. B. germanica colony and tissue sampling

Newly emerged *B. germanica* adult females were obtained from a colony fed *ad libitum* on Panlab 125 dog food and water, maintained in darkness at 29 ± 1°C and 60-70% relative humidity (Belles et al., 1987). All dissections were performed on CO_2_-anesthetized specimens. In adult females, the length of the BOF was used to establish physiological age. Females were maintained with males, and at the end of each experiment, the presence of spermatozoa in the spermatheca was assessed to confirm mating and sperm transfer.

### 2.2. Small RNA library processing

*B. germanica* piRNA sequences were obtained from small RNA libraries previously prepared in our laboratory (Ylla et al., 2017), which are publicly available at GEO, under accession number GSE87031. These sequences correspond to various developmental stages, including ovaries from 7-day-old adults, non-fertilized eggs, embryos (days 0, 1, 2, 6, and 13), and nymphs (instars 1, 3, 5, and 6). The small RNA libraries were preprocessed to remove sequencing adapters using Trimmomatic (v 0.39; relevant parameters used were: ILLUMINACLIP TruSeq2-PE.fa:2:15:10 LEADING:20 MINLEN:18) (Bolger et al., 2014). Reads ranging from 26 to 31 nucleotides were selected using Cutadapt (v 3.5) (Martin, 2011) and aligned to the *B. germanica* genome assembly (v 1.1) (Harrison et al., 2018) with Bowtie2 (v 2.4.4; relevant parameters used were: -a --end-to-end --score-min L,0,0) (Langmead and Salzberg, 2012), resulting in 2,534,205 aligned putative piRNA sequences. Infrequent sequences were filtered out by applying a cut-off of 17 reads across the entire dataset, reducing the number of sequences to 128,653. Finally, the sequences that shared the same genomic 5’ start position and only differed in their length at the 3’ end were collapsed. The counts of each of the collapsed variants were attributed to the most expressed one and normalized using the median-of-ratios method from DESeq2 (Love et al., 2014). The resulting data were used to select the piRNA candidate for functional studies.

### 2.3. Treatments with antisense oligonucleotides

The piRNA-305221 levels were reduced using a chemically unmodified antisense oligonucleotide (ASO: 5’-GGAGGTCCCCCAGACCGGCACAGACCGAA-3’) designed to encompass the entire piRNA sequence. Newly emerged adult females were treated with 1 μL of a solution containing 15 μg/μL of ASO in water (hereafter referred to as ASO-treated), injected in the ventral part of the abdomen with a Hamilton ® 75N syringe. The same dose of an unspecific custom-designed oligonucleotide (Mock) was administered as a negative control. The sequence of Mock corresponds to the concatenated recognition sites of four restriction enzymes (XbaI, HindIII, KpnI, BamHI), and four additional nucleotides to reach the length of the ASO (5’-TCTAGAAAGCTTGGTACCGGATCCCAGGT-3’). Alternatively, both water and Mock were used in control animals since no significant differences were observed in oviposition rates or the number of emerged nymphs (Mock: 40.27 ± 1.161 nymphs, n = 15; Water: 42.25 ± 0.75 nymphs, n = 12; p-value: 0.187), henceforth referred to as Mock-treated.

### 2.4. RNA extraction and expression studies

Ovarian small RNAs were extracted using the miRNAeasy Mini Kit (Qiagen), following the manufacturer’s protocol. The quantity and quality of the extracted small RNAs were estimated by spectrophotometric absorption at 260/280 nm using a Nanodrop spectrophotometer (MicroDigital Co, Ltd). A total of 400 ng from each RNA extraction was reverse transcribed with the 1st Strand cDNA synthesis Kit (Agilent Technologies) to obtain cDNA. The forward primer used in qRT-PCR was designed using the full piRNA sequence, while the universal primer from the Agilent 1st Strand cDNA synthesis Kit was used as the reverse primer. The efficiency of the primers used in qRT-PCR was first validated by constructing a standard curve based on three serial dilutions of cDNA from ovaries (Figure S1). The *U6* small nuclear RNA was used as a reference for expression studies (Tanaka and Piulachs, 2012). qRT-PCR reactions were performed with iTaq Universal SYBR Green Supermix (Bio-Rad Laboratories). Amplification reactions were performed at 95°C for 5 min, 44 cycles of 95°C for 10 s plus 62°C for 40 s, followed by 95°C for 1 min and finally, the melting curve: from 60°C to 95°C with a measurement every 0.5°C increase. After the amplification phase, a dissociation curve was carried out to ensure the presence of only a single product in the amplification (Figure S1).

Total RNA extraction from 7-day-old adult *B. germanica* ovaries was performed using the Tissue Total RNA Purification Kit (Canvax Biotech), following the manufacturer’s protocol. The quantity and quality of the extracted RNAs were estimated by spectrophotometric absorption at 260/280 nm using a Nanodrop spectrophotometer (MicroDigital Co, Ltd). A total of 200 ng of each RNA extraction was reverse-transcribed using the Transcriptor First Strand cDNA Synthesis Kit (Roche) to obtain the corresponding cDNAs. The expression of selected mRNAs related to the studied processes was determined by qRT-PCR, using iTaq Universal SYBR Green Supermix (Bio-Rad Laboratories), and *actin-5c* mRNA expression as a reference. The amplification reactions were performed at 95°C for 3 min, 44 cycles of 95°C for 10 s plus 57°C for 1 min, followed by 95°C for 10 s, and finally, the melting curve: from 57°C to 95°C with a measurement at each 0.5°C increase.

Expression levels of small RNAs and mRNAs were calculated relative to their respective reference gene, using the 2-ΔCt method (Irles et al., 2009b; Livak and Schmittgen, 2001). Results are given as copies of small RNA per 1000 copies of *U6* or copies of mRNA per 1000 copies of *actin-5c* mRNA and correspond to three biological replicates. The primer sequences are detailed in Table S1.

### 2.5. Ecdysone measurements

Ovaries of 7-day-old adults were stored individually at −20°C in methanol to measure ecdysteroids (ecdysone and 20-hydroxyecsdysone). Ecdysteroids were measured by HPLC MS/MS analysis as previously described (Marešová et al., 2024). In brief, after methanolic extraction, detectability of the ecdysteroids is increased 16-to 20-fold by conversion to their 14,15-anhydrooximes. These are further purified by pipette tip solid-phase extraction on a three-layer sorbent and subjected to HPLC-MS/MS analysis (Marešová et al., 2024).

### 2.6. Microscopy methodologies

#### 2.6.1. In situ *hybridization*

The piRNA-305221 was localized in ovaries from *B. germanica* adults using an antisense LNA (locked nucleic acid) probe conjugated to Digoxigenin (DIG) at the 5’ and 3’ ends (5’-DIG-GGAGGTCCCCCAGACCGGCACAGACCGAA-DIG-3’, Merck). Ovaries were dissected in Ringer’s saline and immediately fixed in paraformaldehyde (4% in PBS 0.2 M; pH 6.8) overnight. Subsequent hybridization and washing reactions were carried out as previously reported (Irles et al., 2009b). The DIG hapten was detected by incubating the samples in a 1:4.000 dilution of anti-DIG-rhodamine antibody (Roche, Basel, Switzerland) in PBS-T for 90 min at room temperature. Ovaries were again washed and incubated in DAPI (4’,6-diamidino-2-phenylindole 1 μg/mL, Merck) for 5 min at room temperature. Samples were mounted with Mowiol (Calbiochem) and analyzed by epifluorescence using a Zeiss Axiolmager.Z1 microscope (Apotome) (Carl Zeiss MicroImaging).

#### 2.6.2. DAPI-TRITC-Phalloidin staining

Ovaries from 7-day-old *B. germanica* adults were dissected under Ringer’s saline and immediately fixed in paraformaldehyde (4% in PBS 0.2 M; pH 6.8) for two hours. Subsequently, they were washed with PBT (PBS 0.2 M; pH 6.8 + 0.2% Tween-20) (Irles and Piulachs, 2014). The ovaries were incubated for 20 min in 300 ng/mL of TRITC-phalloidin (tetramethyl rhodamine isocyanate-phalloidin, Merck) prepared in PBT, and after washing with PBT, were incubated for 5 min in 1 μg/mL of DAPI, and washed again with PBT. Samples were mounted with Mowiol (Calbiochem) and analyzed by epifluorescence using a Zeiss Axiolmager.Z1 microscope (Apotome) (Carl Zeiss MicroImaging).

#### 2.6.3. Embryo observations

Fifteen-day-old oothecae were removed from the female abdomen by gentle pressure to observe the embryo development. The oothecae were incubated for 5 min in water at 95°C to facilitate the individualization of the embryos. Images of the embryos were obtained using a Zeiss DiscoveryV8 stereomicroscope (Carl Zeiss MicroImaging).

### 2.7. Statistical analysis

The data are expressed as mean ± standard error of the mean (S.E.M.). Statistical analyses were performed using GraphPad Prism version 8.1.0 for Windows, GraphPad Software. Significant differences between control and treated groups were calculated using the Student’s t-test. Data were evaluated for normality and homogeneity of variance using the Shapiro–Wilk test, which indicated no transformations were needed. All datasets passed the normality test.

## Results

### 3.1. piRNA-305221 in *Blattella germanica* ovary

*B. germanica* piRNAs were ranked according to their levels in the libraries of small noncoding RNAs of 7-day-old adult ovaries and non-fertilized eggs, obtained previously (Ylla et al., 2017). One of the most abundant piRNAs in ovaries of 7-day-old adults, and with quite high levels in non-fertilized eggs, is the piRNA-305221 (5’-UUCGGUCUGUGCCGGUCUGGGGGACCUCC-3’) (Figure 1A), suggesting that it is maternally provided. The piRNA-305221 levels are still detectable in early embryonic stages (days 0 to 2), becoming undetectable in later embryo development and early nymphal instars. It reappears, albeit at low levels, in the last nymphal instars (Figure 1A).

**Figure 1.**
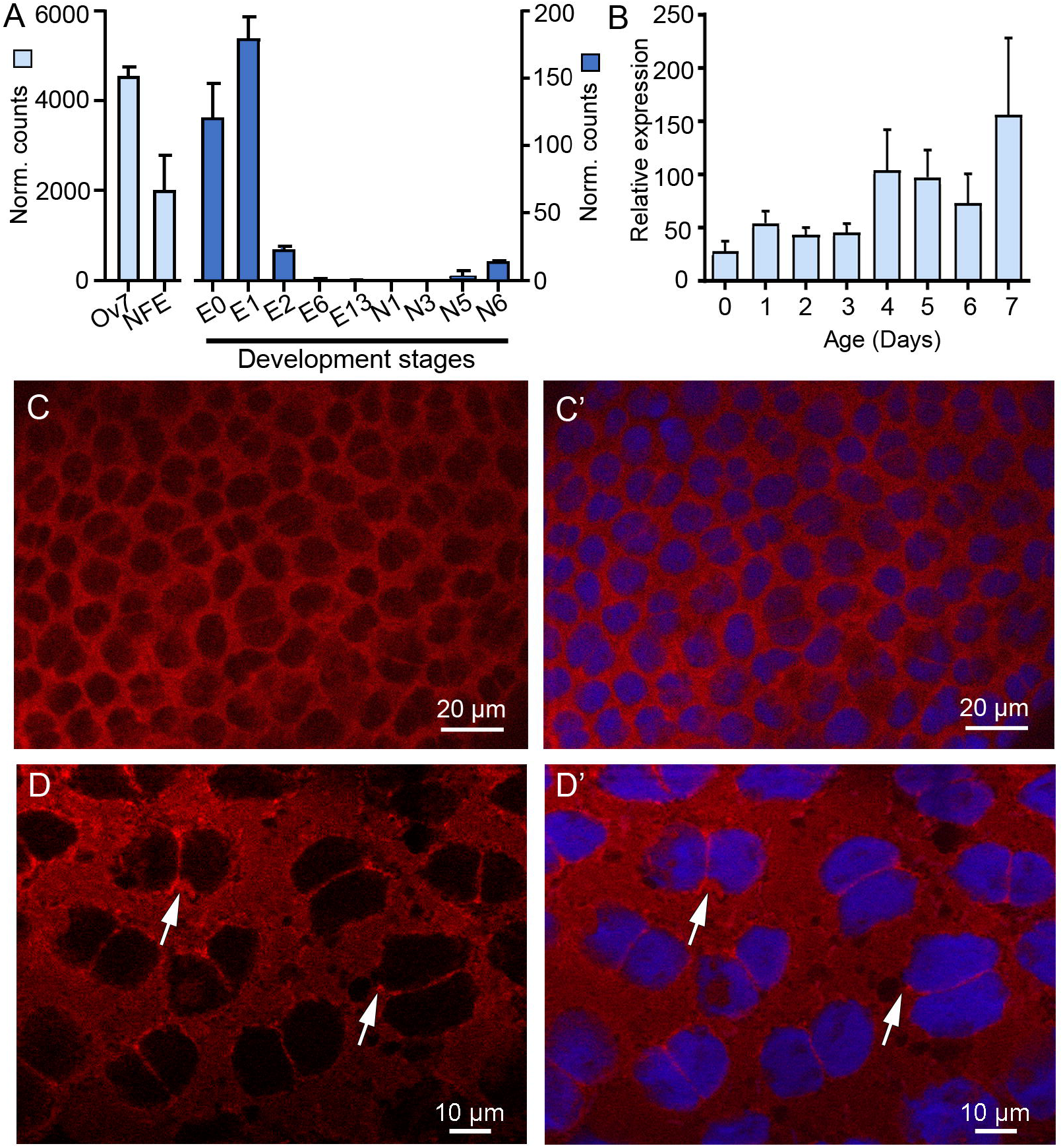
piRNA-305221 in *Blattella germanica* adult ovaries. **A**. Levels of piRNA-305221 in the small RNA libraries from different developmental stages, including embryos of different ages (days 0, 1, 2, 6 and 13 (E0, E1, E2, E6, E13)), nymphs from first (N1), third (N3), fifth (N5) and sixth (N6) instars. Data from transcriptomes from 7-day-old adult ovaries (Ov7) and non-fertilized eggs (NFE) were also included (Ylla et al., 2017). Each value represents the average of normalized counts, conducted using the median-of-ratios method from DESeq2. Data are shown as mean ± S.E.M. (n = 2). **B**. Expression profile of piRNA-305221 in adult ovaries during the first gonadotropic cycle. Data represent copies of piRNA per 1000 copies of *U6*, and are expressed as the mean ± S.E.M. (n = 3). **C**. *In situ* hybridization in late 3-day-old adult ovaries showing the piRNA-305221 localization in the cytoplasm of follicular cells of a BOF. **C’** shows the merged images with the nuclei stained with DAPI. **D**. *In situ* hybridization in follicular cells from 5-day-old adult ovaries. **D’**. shows the merged images with the nuclei stained with DAPI. piRNA-305221 labeling accumulates close to the nuclear membrane (arrows). An antisense LNA probe with the piRNA sequence was used and labeled with DIG at both ends. In C-D, the probe was revealed with a rhodamine-labeled anti-DIG antibody (red). The DNA in the nuclei was stained with DAPI (blue).

The expression of the piRNA-305221 was measured by real-time PCR in *B. germanica* adult ovaries. Twenty-four hours after the adult emergence, its expression increases with a fold-change close to two, and these levels are maintained until adult day-3. In the transition from 3-to 4-day-old females, the expression increases twofold again (Figure 1B), coinciding with important changes in the BOF. In 3-day-old adults, the FCs arrest cytokinesis, becoming binucleated and beginning to be polyploid (Irles and Piulachs, 2014). From day 4, piRNA-305221 levels decrease smoothly, rising again on day 7 when it reaches its highest levels (Figure 1B), concurrently with the synthesis of the chorion proteins by the FCs (Irles et al., 2009b). *In situ* hybridization revealed the piRNA-305221 spread in the cytoplasm of FCs in the BOF of adult females (Figure 1C and D). In mature BOF, the piRNA-305221 label is more intense near the nuclear membrane (Figure 1D and D’, arrows), where piRNAs are normally found.

### 3.2. Function of piRNA-305221 in *Blattella germanica* ovary

To study the piRNA-305221 function, newly emerged *B. germanica* adult females were treated with an ASO targeting the piRNA, and the phenotypes were observed 7 days after the treatment.

In ovaries of 7-day-old treated females, levels of the piRNA-305221 showed a significant reduction, close to 50% (p = 0.0107) compared to Mock females (Figure 2A). This depletion results in a delay in BOF growth. The length of the BOFs in the ASO-treated females was significantly smaller (1.697 ± 0.05 mm; n = 31; p < 0.0001) than in Mock females (2.037 ± 0.03 mm; n = 11) (Figure 2B), although the variability in treated individuals was rather wide. In 7-day-old Mock adult females, the binucleated FCs in the BOF exhibit uniform distribution, beginning to narrow the intercellular spaces (Figure 2C, arrows), with nuclei of the same size and shape (Figure 2D). In contrast, in ASO-treated females, the FCs in the BOF exhibited enlarged nuclei with heterogeneous shapes and bizarre morphologies (Figure 2E).

**Figure 2.**
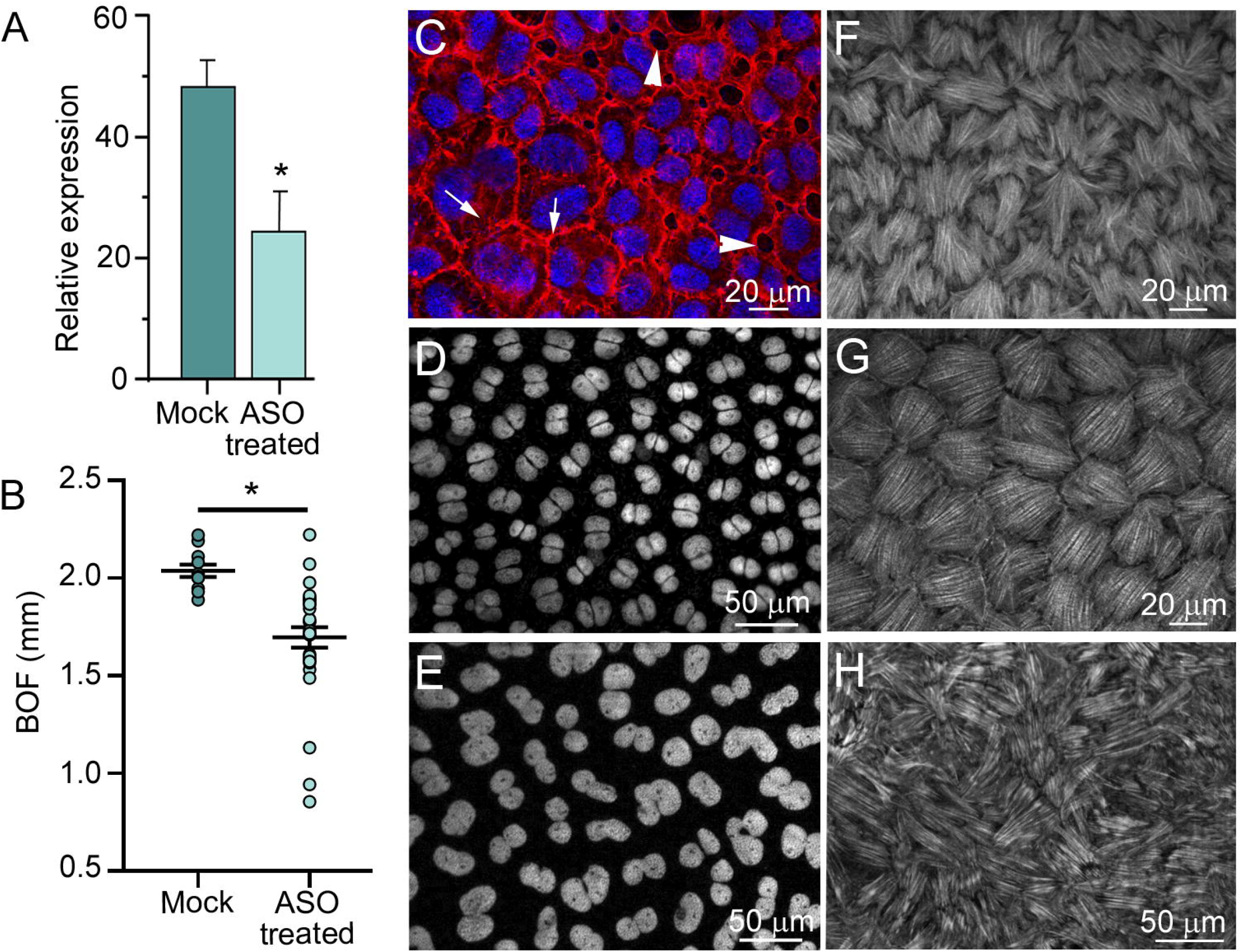
Function of piRNA-305221 in ovaries. Newly emerged *B. germanica* adult females were treated with ASO or Mock at emergence and dissected 7 days later. Images from C-H correspond to basal ovarian follicles. **A**. Expression levels of piRNA-305221 in 7-day-old adult Mock or ASO-treated ovaries. Data represent copies of the piRNA per 1000 copies of *U6*, and are expressed as the mean ± S.E.M. (n = 3). The asterisk indicates statistically significant differences (p = 0.01). **B**. Basal ovarian follicles (BOFs) lengths (mm) in Mock (n = 11) and ASO-treated (n = 31) females, measured in 7-day-old adults. Data were represented as mean ± S.E.M. The asterisk indicates statistically significant changes (p < 0.0001). **C**. Follicular epithelia, from a Mock female, showing the follicular cells (FCs) distribution; cells are closing the intercellular spaces; arrows indicate closed intercellular spaces, and arrowheads indicate intercellular spaces that are still open. **D**. Nuclei from FCs in Mock female, showing the uniformity in size and distribution. **E**. Nuclei from FCs in ASO-treated females, showing different sizes and forms. **F**. F-actin distribution in the basal pole of FCs from a Mock female in early chorion stage; the actin fibers in the basal pole of FCs are not completely packed. **G**. F-actin distribution in the basal pole of FCs from a Mock female in late chorion stage; the F-actin fibers are completely packed, giving the FCs a round shape. **H**. Disorganization of F-actin distribution in the basal pole of FCs of ASO-treated late 7-day-old females; the F-actin microfilaments were stained with phalloidin-TRITC (red in C, white in F, G, and H). DNA was stained with DAPI (blue in C, white in D and E).

In *B. germanica* ovaries, upon completion of the chorion, the actin cytoskeleton in BOFs surrounds the FCs, modifying their distribution and covering the surface of the basal pole of FC, in an orderly manner (Figure 2F, G). Just before ovulation, the actin fibers wrap up the FCs, giving them a rounded shape (Figure 2G), which facilitates the contractions of the ovarian follicle needed for ovulation and oviposition. However, in ASO-treated females, the distribution of the F-actin fibers in the FCs cells of the basal pole was disorganized (Figure 2H). The fibers did not follow the correct orientation on the ovarian follicle surface and appeared to overlap different FCs.

### 3.3. piRNA-305221 is involved in *Blattella germanica* reproduction

We then investigated whether depletion of piRNA-305221 in adult females could affect oviposition and embryo viability. Newly emerged adult females were treated with the ASO, then they were paired with males, and observed until ootheca formation. The first effect observed was a significant delay in oviposition in ASO-treated (8.11 ± 0.26 days; n = 53; p = 0.013), compared to Mock females (7.33 ± 0.08 days; n = 38).

Despite this delay, ASO-treated females carried the ootheca throughout the entire embryogenesis period (18.50 ± 0.26; n = 32) with no significant differences from Mock females (18.81 ± 0.21; n = 28). However, although all the ASO-treated females successfully mated (n = 32), a significant number (25%, n = 8) of oothecae failed to produce nymphs, and in the remaining ASO-treated females, the number of hatching nymphs was significantly reduced (36.29 ± 1.60; n = 24) compared to Mock females (41.15 ± 0.74; n = 27; p = 0.0062). The lower number of hatching nymphs may be linked to the production of defective oothecae in several ASO-treated females (31.25%, n = 10), with shortened, curved, or dried oothecae, or a combination of these defects, which often resulted in misaligned eggs, compromising viability and hatching (Figure 3A and B). The number of hatching nymphs from these malformed oothecae was dramatically reduced (24.25 ± 3.64 nymphs; n = 4; Figure 3B), and in some cases, no nymphs hatched at all (n = 6).

**Figure 3.**
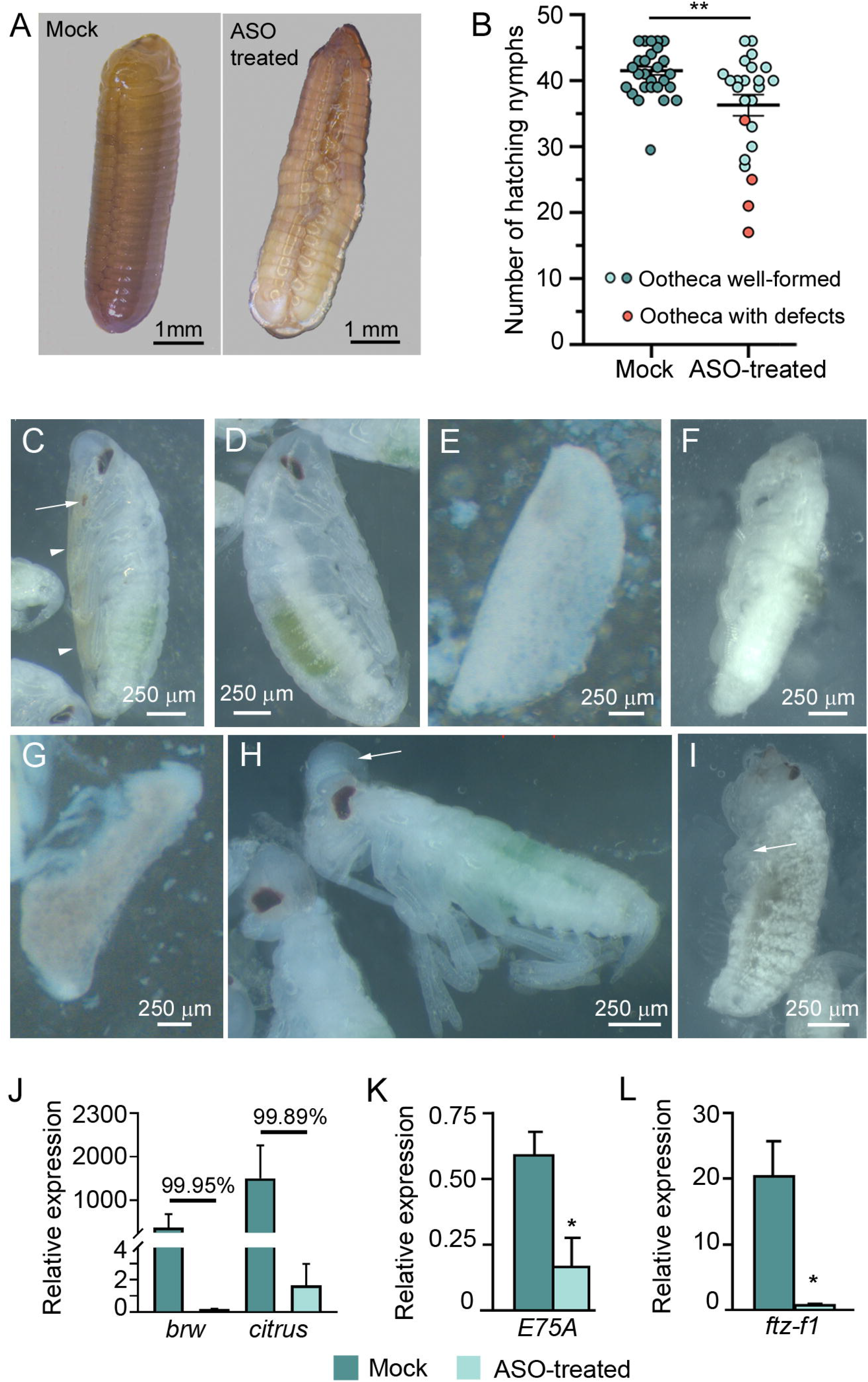
piRNA-305221 on reproduction and embryogenesis. **A**. Fifteen-day-old oothecae from Mock and ASO-treated females; the latter shows the eggs were not well-aligned. **B**. Number of nymphs hatching from oothecae formed by Mock and ASO-treated females. Each dot represents the total nymphs from one ootheca. Blue dots represent well-formed oothecae, and orange dots represent defective oothecae. Data are represented as the mean ± S.E.M. (Treated females, n = 27; Control females, n = 24). Asterisks indicate statistically significant differences (** p = 0.01). **C**. Fifteen-day-old embryos from Mock females. Segmentation is complete; the mandibles (arrow) and legs (arrowheads) sclerotization has already started. **D-I**: Fifteen-day-old embryo from ASO-treated females. **D**. Embryo similar to controls without sclerotized appendages. **E**. Embryo with no signs of development. **F**. Embryo with the different body parts differentiated; the appendages have some degree of development; the eyes are not pigmented. **G**. Similar to F, but less advanced development. The legs are distinguishable, but the body segmentation is not apparent. **H**. Embryo with correct body development; however, the head has an abnormal morphology, and the dorsal organ is still visible (arrow); these modifications determine a mislocalization of the head appendages. In addition, the legs appear to be larger than in controls. **I**. Embryo with complete body segmentation; the head does not have the correct structure; the eyes are correctly pigmented, but the head appendages are not well defined (arrow); the legs are short, reaching only the first abdominal segments; there is an increase in cells corresponding to the fat body that appeared like white dots. **J**. Expression of *brownie* (*brw*) and *citrus*. **K**. Expression of *E75A*. **L**. Expression of *fushi tarazu-f1* (*ftz-f1)*. In J - L, data represent copies of mRNA per 1000 copies of *actin-5c* mRNA of 7-day-old ovaries from adult females treated with Mock or ASO at adult day 0, and are expressed as the mean ± S.E.M. (n = 3-5). The asterisks indicate statistically significant differences in ASO-treated respect to Mock * p = 0.03 (*E75A*); * p = 0.02 (*ftz-f1*).

We examined 15-day-old embryos from Mock and ASO-treated females, an age at which the embryos are nearly fully developed before hatching on day 18. At this stage, embryos from Mock females (a total of 94 embryos from 4 different oothecae) have completed dorsal closure, and the tips of the antennae and hind legs have almost reached the seventh abdominal segment. Eye pigmentation is complete, and the sclerotization of the mandibles and legs is beginning. (Figure 3C) (Piulachs et al., 2010; Tanaka, 1976). In contrast, embryos from ASO-treated females (107 embryos from 3 different oothecae) exhibited a wide range of phenotypic defects. 62% of embryos were morphologically similar to controls but showed less sclerotization (Figure 3D). The phenotypes of the remaining 38% of embryos displayed a variety of defects, ranging from eggs lacking the germinal band or with some tissue concentration in the ventral side, with the aspect of an amorphous mass (Figure 3E), to embryos with incomplete segmentation and with defective eyes (Figure 3F, G). Some embryos lacked eye pigmentation (Figure 3F), while others had eyes of incorrect shape (Figure 3H). The dorsal organ was still visible in this group of embryos (Figure 3H, arrow), and none showed signs of appendage sclerotization, and, in general, the legs were improperly sized and shaped (Figure 3F-I, arrow in I). In addition, in a few ASO-treated embryos, the fat body cells (urate cells and mycetocytes, see A. Tanaka, 1976) were more abundant than in Mock embryos (Figure 3I, white dots in the thorax and abdomen).

The changes in FCs, with the problems in ootheca formation and embryo development, suggest changes at the chorion levels. We quantified two genes involved in chorion synthesis, and the results revealed a dramatic depletion of brownie (*brw*) and *citrus* (99.95% and 99.89%, respectively; Figure 3J), although the differences with respect to the controls were not statistically significant due to the dispersion of the data.

This dispersion likely reflects the restricted temporal and spatial expression of these genes, which are only active in the follicular epithelium during chorion synthesis (Irles and Piulachs, 2011; Irles et al., 2009a). The depletion of *brw* and *citrus* expressions suggests a disruption of ecdysone signaling in the ovary, as was indicated by a significant reduction of *E75A* expression (71%; p = 0.03; Figure 3K) and a dramatic depletion of that of *ftz-f1* (94%; p = 0.02; Figure 3L).

### 3.4. piRNA-305221 modulates the production of 20E

To determine whether piRNA-305221 influences ecdysone synthesis or signaling, we examined the expression of steroidogenic genes in the ovaries of ASO-treated females (Figure 4). We observed significant changes in mRNA levels. While the expression of *nvd* and *sad* remained unaffected, *spo, phm*, and *dib* expression increased significantly by 88.52% (p = 0.0150), 159.93% (p = 0.0105), and 133.19% (p = 0.0145), respectively (Figure 4A). Interestingly, the expression of *shd*, which encodes an ecdysone 20-monooxygenase that mediates the hydroxylation of ecdysone into 20-hydroxyecdysone (20E), decreases significantly by an average of 58.32% (p = 0.0031) (Figure 4B). This reduction in *shd* expression likely affects the availability of the 20E in the ovary, as was suggested by the E75 depletion (Figure 3K). To corroborate this data, we quantified by HPLC MS/MS the ecdysone and 20E content in ovaries of Mock and ASO-treated females (n = 4) (Figure 4C). When comparing the ecdysone levels of controls versus ASO-treated females, we observed a reduction of 56.47% in the ASO-treated group, although the differences were not statistically significant, due to the dispersion in control samples. Conversely, the 20E levels, the active form, showed a dramatic decrease in ASO-treated ovaries, with a reduction by 93.78% compared to controls (Figure 4C). Hormone measurements revealed a substantial hormonal imbalance in ASO-treated females. The relationship between 20E and ecdysone in Mock ovaries is 0.689, indicating an equilibrium between the ecdysone synthesized and its transformation to 20E. Conversely, in ASO-treated females, the relationship between 20E and ecdysone is 0.099, showing the imbalance between both, with ecdysone accumulated in the ovary. All these data indicate that piRNA-305221 is involved in ecdysteroid biosynthesis, promoting the conversion of ecdysone into 20E in the ovary of *B. germanica*.

**Figure 4.**
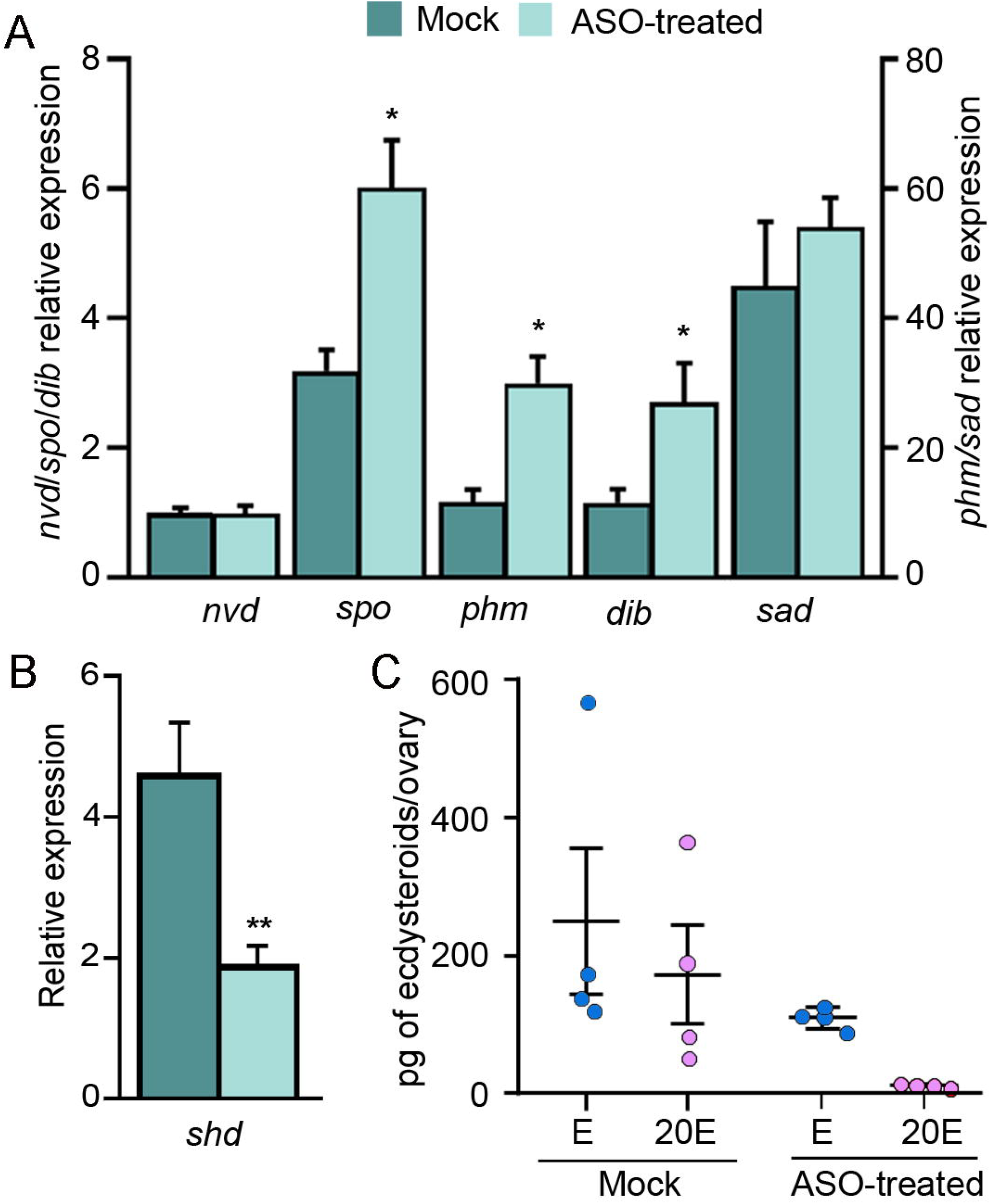
piRNA-305221 depletion affects steroidogenic genes. **A**. Expression of the steroidogenic genes, *neverland* (*nvd*), *spook* (*spo*), *phantom* (*phm*), *disembodied* (*dib*), and *shadow* (*sad*). **B**. Expression of shade (*shd*). **C**. Scatter plot showing the relationship between ecdysone (blue dots) and 20-hydroxyecdysone levels (pink dots) in ovaries of Mock and ASO-treated females. In A and B, data represent copies of mRNA per 1000 copies of *actin-5c* mRNA of 7-day-old ovaries from adult females treated with Mock or ASO at adult day 0, and are expressed as the mean ± S.E.M. (n = 3-6). The asterisks indicate statistically significant differences respect to Mock: * p = 0.0150 (*spo*); * p = 0.0105 (*phm*); * p = 0.0145 (*dib*): ** p = 0.0031 (*shd*). In C, data represent picograms (pg) of ecdysteroids in a single 7-day-old ovary of each female (each dot) from adult females treated with Mock or ASO at adult day 0, and are expressed as the mean ± S.E.M. (n = 4).

## 4. Discussion

The role of piRNAs as regulators of TEs and genome protectors is well-established. However, recent research has shown the importance of piRNAs as mRNA regulators. Thus, piRNA functions have been extended to processes as different as the maintenance of germinal stem cells, DNA repair mechanisms, sex determination, chromatin modifications, learning and memory, and cancer mechanisms (Gleason et al., 2018; Iki et al., 2023; Kiuchi et al., 2014, 2023).

Insect oogenesis is a highly regulated event involving multiple steps and processes, including germ cell proliferation, oocyte growth and maturation, vitellogenesis, and eggshell formation. These processes occur under the control of ecdysteroids and juvenile hormone (Alborzi and Piulachs, 2023; Belles et al., 2024). Due to the efficiency and precision required, the regulation of all these processes entails different regulatory layers involving proteins, mRNAs, and non-coding RNAs, which have specific functions at different regulatory levels. Among the non-coding RNAs are the small non-coding RNAs that include the piRNAs studied here.

The piRNA-305221 is highly expressed in *B. germanica* ovaries, and is present in somatic and germinal cells, as is evidenced by its presence in unfertilized eggs (Llonga et al., 2018). Furthermore, piRNA-305221 expression in adult ovaries correlates with key changes in the BOF, which are mainly related to changes in the FCs program (Claycomb and Orr-Weaver, 2005; Irles et al., 2016, 2009b; Irles and Piulachs, 2014, 2011). The changes in piRNA-305221 expression during critical moments, such as the arrest of cytokinesis, the increase of polyploidy, or endoreplication in the FCs, suggest a role in genome protection. However, piRNA-305221 is maternally provided to the embryo, suggesting further functions.

Depletion of piRNA-305221 using an ASO enabled us to infer its functions in the *B. germanica* adult ovary. Thus, decreased levels of piRNA-305221 in the ovary determined a delay in oviposition and malformations in the ootheca shape and size. These oothecae defects could explain the observed reduction in emerging nymphs. Moreover, the expression profile of the piRNA-305221 in the ovary and the observed oothecae malformations after reducing its levels, suggest an influence of this piRNA on genes crucial at the end of the gonadotropic cycle, among them, steroidogenic genes expressed in the adult ovary (Ramos et al., 2020; Belles et al., 2024). In most adult insects, when the prothoracic gland degenerates, the ovary becomes the main source of ecdysone, acting in an autocrine manner (Pascual et al., 1992; Romaña et al., 1995). In *B. germanica* adult females, ecdysone is primarily needed at the end of the gonadotropic cycle to facilitate the endoreplication of some genome regions in the FCs, promoting chorion synthesis and facilitating the egg release into the oviduct (Alborzi and Piulachs, 2023; Belles et al., 2024; Belles et al., 1993). Depletion of piRNA-305221 led to reduced *shd* mRNA expression, which prevents the conversion of ecdysone to 20E, and resulted in an overexpression of *spo, phm*, and *dib*. Since the piRNAs have been described as repressors of mRNAs, the reduction of *shd* expression observed after the ASO treatment suggests that piRNA-305221 would inhibit a *shd* repressor. Concomitantly, the overexpression of *spo, phm*, and *dib* might be explained by a feedback mechanism triggered by the interruption of the 20E flow.

The diversity of phenotypes resulting from piRNA depletion could be attributed to the piRNA-305221 action on different mRNA targets. It is worth noting that the functions described here may represent only part of the roles of piRNA-305221 in *B. germanica* ovaries. The maternal provision of piRNA-305221 to the embryo suggests that it would play specific roles in early embryo development, a topic for future research. Nonetheless, our results show that ASOs-mediated piRNA depletion can be a useful approach to study the functions of these still mysterious sncRNAs.

## Acknowledgments

We thank the financial support to the project PID2021-122316OB-I00 from the MCIN/AEI/ 10.13039/501100011033 and by ERDF, a way of making Europe, project 25-154535 from the Czech Science Foundation, Czech Republic, to MN. This work was also supported by Pla de Doctorats Industrials de la Secretaria d’Universitats i Recerca del Departament d’Empresa i Coneixement de la Generalitat de Catalunya (grant number 2021 DI 059), and AGAUR 2021 SGR 00419). The authors acknowledge the support of the publication fee by CSIC Open Access Publication Support Initiative through its Unit of Information Resources for Research (URICI).

