## Supplementary figures and images for "A piRNA modulates the levels of 20-hydroxyecdysone in the ovary of the German cockroach"

### Figure S1

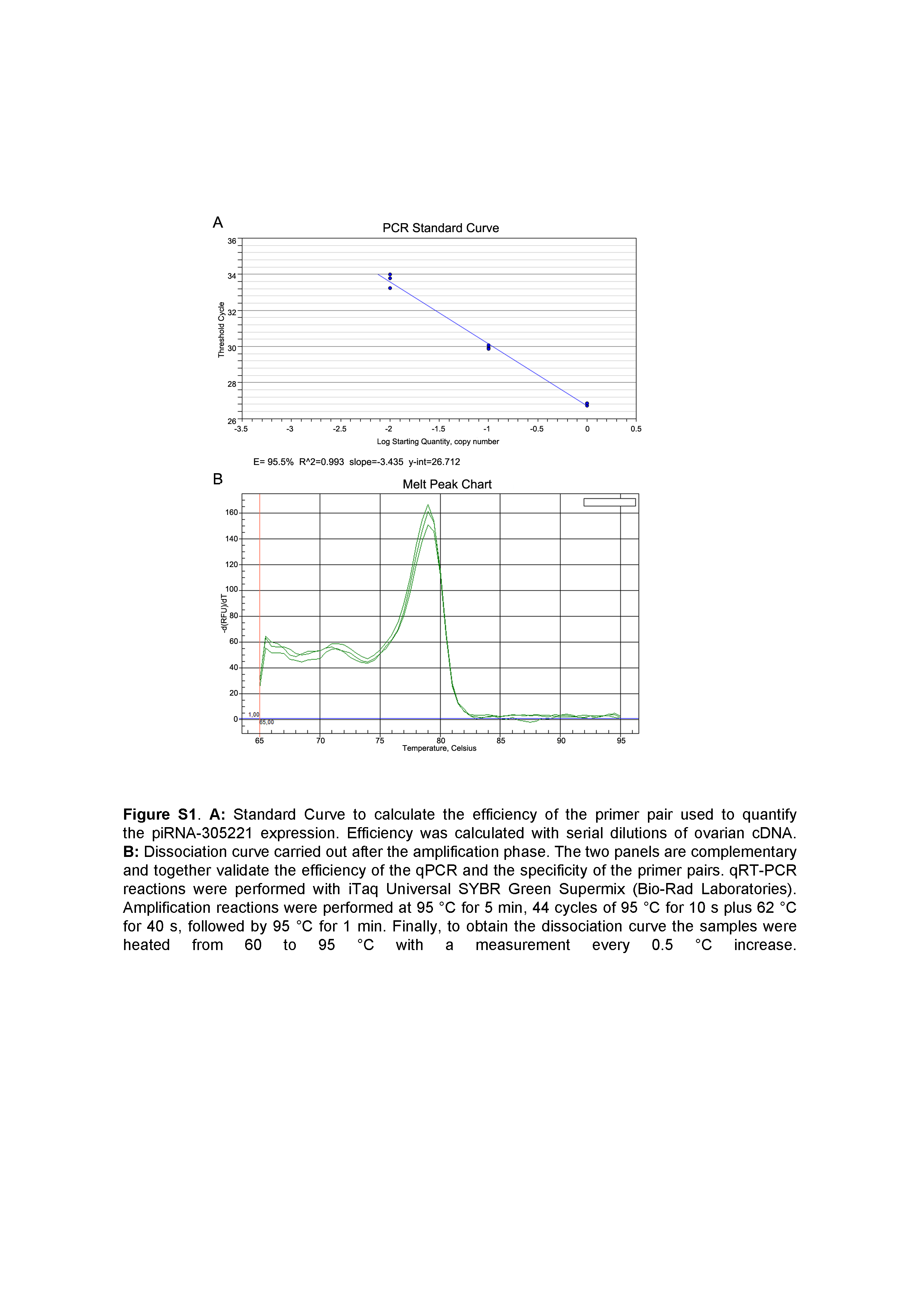
