## Supplementary material for "A piRNA modulates the levels of 20-hydroxyecdysone in the ovary of the German cockroach": Table S1

**Supplementary Table 1.** Primers used in the qRT-PCR. F: forward primer, R: reverse primer. The corresponding accession number in NCBI is included.

| **Name** |  | **Primer sequence (5’-->3’)** | **Accession number** |
| --- | --- | --- | --- |
| piRNA_305221 | F | TTCGGTCTGTGCCGGTCTGGGGGACCTCC | - |
| Universal (Agilent) | R | GACGAGCTGCCTCAGTCGCATA | - |
| U6 | F  R | CGATACAGAGAAGATTAGCATGG  GTGGAACGCTTCACGATTTT | FR823379 |
| *actin-5C* | F  R | AGCTTCCTGATGGTCAGGTGA  TGTCGGCAATTCCAGGGTACATGGT | AJ862721 |
| *dib* | F  R | GCAACAGACAATGGACCTCA  AGATCCAATGCAACCTCCTC | PSN36324 |
| *E75A* | F  R | AATGAGTAGAGATGCGGTGCGGTT  TCAGCGTCGGACAGTCTTAGTGA | CAJ87513.1 |
| *ftz-f1* | F  R | TTGTCACATCGACAAGACGCA  GTACATCGGGCCGAATTTGTTTCT | FM163377 |
| *nvd* | F  R | CTGGGGCCAGTCACAATACT  GCAGGGGCTTGTCAATGTAT | PSN31862 |
| *phm* | F  R | CTAGGCACCAGAGCACCTTC  GCAAGCACTGTGTCTTCCAA | PSN36025.1 |
| *sad* | F  R | ATGAGGAGGTTCAGGGTGTG  CTGGCCAGAAGTCATTTGGT | OE845190.1 |
| *shd* | F  R | CACAGAGGCGCACAAGTTTA  GTTCCCCTTCAAAGTCCACA | PSN43891.1 |
| *spo* | F  R | GCCTTCATCATGTTGGCGTC  CAGGTGTGGAGAGGTGTCTG | PSN30774.1 |
| *citrus* | F  R | TCGTGCTTTTCAATGTGCGTA  GGGAATCCAGGGTATTTGGAA | FN823078.1 |
| *brownie* | F  R | CTCAGCACAAAGCCGTAGCA  CGTCGGCGTAAGCTTCGTAG | FM253364.1 |
